# Deep brain stimulation pulse sequences to optimally modulate frequency-specific neural activity

**DOI:** 10.1101/2022.03.12.484100

**Authors:** Hafsa Farooqi, Jerrold L. Vitek, David Escobar Sanabria

## Abstract

Precise neuromodulation systems are needed to identify the role of neural oscillatory dynamics in brain function, and to advance the development of electrical stimulation therapies tailored to each patient’s signature of brain dysfunction. Low-frequency, local field potentials (LFPs) are of increasing interest for the development of these systems because they reflect the synaptic inputs to a recorded neuronal population and can be chronically recorded in humans. Here, we identified stimulation pulse sequences to optimally minimize or maximize the 2-norm of frequency-specific LFP oscillations using a generalized mathematical model of spontaneous and stimulation-evoked LFP activity, and a subject-specific model of neural dynamics in the pallidum of a Parkinson’s disease patient. We leveraged convex and mixed-integer optimization tools to identify the pulse patterns, and employed constraints on the pulse frequency and amplitude that are required to keep electrical stimulation within its safety envelope. Our analysis revealed that a combination of phase, amplitude, and frequency pulse modulation is needed to attain optimal suppression or amplification of the targeted oscillations. Phase modulation is sufficient to modulate oscillations with a constant amplitude envelope. To attain optimal modulation for oscillations with a time-varying envelope, a trade-off between frequency and amplitude pulse modulation is needed. The optimized pulse sequences identified here were invariant to changes in the dynamics of stimulation-evoked neural activity, including the damping and natural frequency or complexity (i.e., generalized vs. patient-specific model). Our results reveal the structure of pulse sequences that can be used in closed-loop brain stimulation devices to control neural activity in real-time.

**Significance Statement:** This computational study reveals the structure of electrical pulse sequences that optimally modulate spontaneous, synaptic-related neural oscillations measured using local field potentials (LFPs). We considered constraints on the pulse amplitude and frequency, relevant to keep stimulation pulses within the safety envelope. Our mathematical analysis indicate that these optimized pulse sequences are invariant to changes in the studied neural dynamics, and a trade-off between amplitude and frequency pulse modulation is critical when the pulse amplitude is constrained. The pulse sequences identified here provide the rationale for the development of closed-loop neuromodulation systems that employ stimulation-evoked responses to control neural activity. These closed-loop systems will be key to characterize the role of neural activity in brain function and ultimately advance personalized neuromodulation devices.

## 1. Introduction

Robust and precise neural control methodologies are needed to characterize the role of neural activity in the manifestation of brain conditions, and to advance the development of neuromodulation technologies. Of particular interest for the development of these technologies are the local field potentials (LFPs) recorded chronically in humans using macro-electrodes given the stability of these LFPs over time (1, 2). Experimental and theoretical studies have shown that LFP neural activity at low-frequency (<100 Hz) is dominated by synchronized synaptic inputs to neuronal populations near the recording site (3, 4). Therefore, these LFPs can be used as a feedback signal in closed-loop systems to control synchronized synaptic activity in a neuronal population and thereby modulate information flowing into and out of the targeted neurons.

In Parkinson’s disease, synchronized 11-35 Hz (“beta” band) LFP oscillations in the basal ganglia are hypothesized to be associated with motor dysfunction (5–10). A recent study with the 1-methyl-4-phenyl-1,2,3,6-tetrahydropyridine (MPTP) nonhuman primate model of Parkinson’s disease showed that amplification or suppression of beta band oscillations in the subthalamic nucleus (STN) could be achieved using STN neural responses evoked by electrical stimulation in the internal segment of the globus pallidus (GPi) (11). The rationale behind this approach, referred to as closed-loop evoked interference deep brain stimulation (eiDBS), is that synaptic-related neural responses evoked by electrical pulses can “override” spontaneous, synaptic-related oscillations via synaptic summation when the pulses are delivered with adequate amplitude and precise timing relative to the targeted oscillation’s phase (11). More recently, the feasibility of using eiDBS to suppress or amplify neural activity (16-22 Hz) in the human GPi using a single deep brain stimulation (DBS) electrode array was demonstrated in a patient with Parkinson’s disease (12). The current implementation of eiDBS delivers electrical pulses of fixed amplitude locked to the phase of frequency-specific oscillations. This approach can be effective to suppress or amplify the mean amplitude of targeted neural activity, but it is sub-optimal to suppress neural oscillations because it does not account for dynamic changes in the amplitude of spontaneous neural oscillations. Because the stimulation pulse amplitude is set constant, variations in the oscillation amplitude can result in unwanted amplification when the spontaneous oscillations are small, or sub-optimal suppression when the spontaneous oscillations are large relative to the stimulation-evoked oscillations. Identifying electrical stimulation strategies to optimally modulate neural activity is needed to address the shortcomings of the present eiDBS implementation and advance the development of precise neuromodulation devices.

In this study, we identified pulse modulation patterns that optimally suppress or amplify the 2-norm of spontaneous neural activity. To discover these pulse patterns, we employed optimization tools together with generalized mathematical models that describe the temporal dynamics of spontaneous and stimulation-evoked neural activity. We considered evoked response dynamics with distinct damping coefficients and natural frequencies to understand whether the structure of the optimal pulse sequences remained invariant with changes in these dynamics. We also derived optimal stimulation sequences using a model of neural activity in the GPi of a Parkinson’s disease patient to confirm that the structure of these pulse sequences is preserved in neural dynamics more complex than those described by the generalized mathematical model. Our methodology enabled us to consider constraints on the pulse amplitudes and frequencies to comply with devices currently cleared or approved by the U.S. Food and Drug Administration (FDA).

This study shows that a trade-off between phase, frequency, and amplitude pulse modulation is needed to optimally minimize or maximize the 2-norm of targeted neural activity. The pulse modulation strategies identified in this study provide the rationale to optimize closed-loop brain stimulation systems and control neural activity in real-time effectively. Optimizing these systems could, in turn, enable us to characterize the role of oscillatory dynamics in brain function and advance the development of precise, personalized neuromodulation technology.

## 2. Methods

### A. Mathematical models of neural dynamics

We employed mathematical models of spontaneous and stimulation-evoked neural dynamics to identify pulse sequences that minimize the 2-norm of frequency-specific neural activity. We considered a generalized mathematical model in which relevant parameters of the stimulation evoked responses (i.e., damping and natural/resonant frequency) were varied in order to understand whether the identified pulses patterns can be applied to distinct neural dynamics. To understand whether our results from the generalized model could be used to modulate more complex neural dynamics similar to those observed in-vivo, we identified optimal pulse sequences for a mathematical model of neural activity in the GPi derived with data recorded from a PD patient (12). We considered electrical stimulation waveforms that consist of a negative followed by a positive square pulse. These two square pulses have the same amplitude and pulse width (i.e., symmetric, biphasic waveform).

The effect of stimulation pulses on the neural evoked response temporal dynamics was characterized using a saturation nonlinearity (static) connected to a system of linear equations of differences (discrete time approximation of differential equations) as depicted in Fig. 1. This model structure has been shown to accurately approximate the linear response of neural circuitry to electrical stimulation pulses (11, 12). The evoked response is described by the following equations of differences.

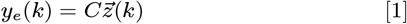

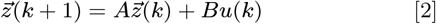

where *k* = 0, 1, …, *n* – 1 is the discrete time sample associated with time *kt_s_, t_s_* is the sampling time, 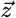 is the state vector, *u*(*k*) > 0 is the saturated input stimulus, *y_e_*(*k*) is the evoked response at sample *k*, and A, B, and C are constant matrices that parameterize the equations of differences. The saturated stimulation input is given by 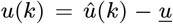 with 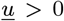 being the minimum stimulation current that evokes a neural response, and 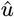 the biphasic stimulation pulse.

**Fig. 1.**
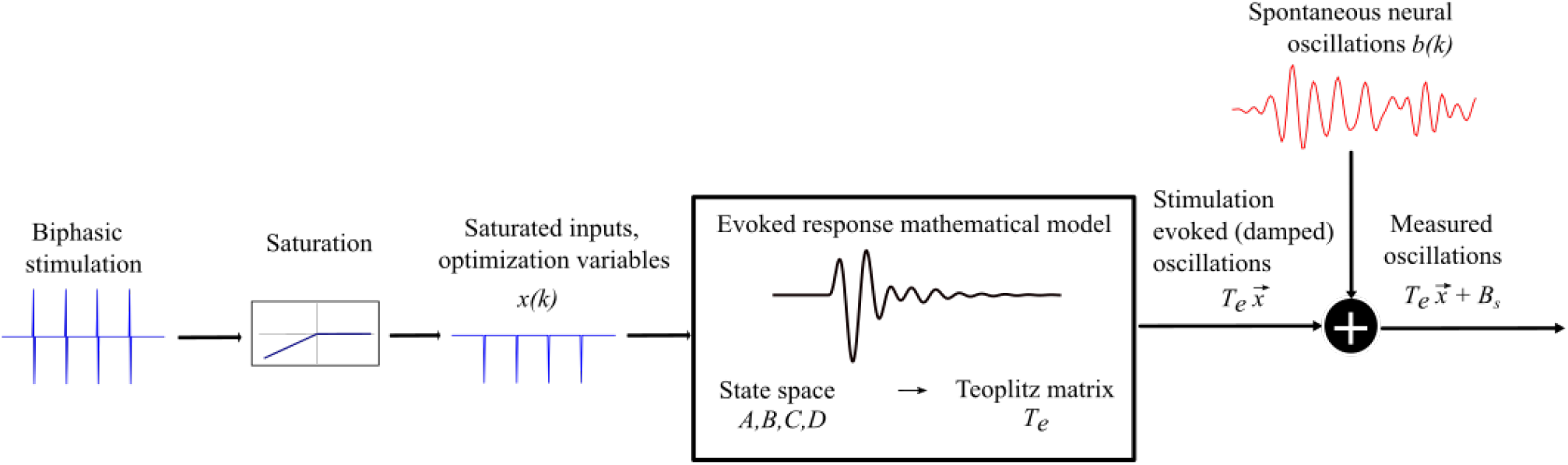
A schematic of the model dynamics capturing the interaction between spontaneous and stimulation-evoked neural activity in the targeted brain region. A biphasic stimulation pulse train, where each pulse is represented by a positive followed by a negative square pulse, which after saturation is reduced to just the negative square pulse train. The saturated pulse train is responsible for evoking stimulation based oscillations, the dynamics of which can be mapped to a teoplitz matrix (*T_e_*). The final model is the summation of the matrix multiplication of the *T_e_* with the optimization variable vector 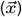 and the vector containing the spontaneous oscillations (*B_s_*).

The LFP measurement, which reflects the combined spontaneous and stimulation-evoked neural activity, is modeled as

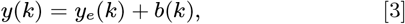

where *b*(*k*) is the value of the spontaneous neural oscillations at sample *k*. These spontaneous oscillations were modeled using sinusoidal functions or synthetic LFP signals that match the power spectral density (PSD) of LFPs recorded from a PD patient.

#### Generalized, low-order model

We used a second-order model of the evoked response to characterize optimal stimulation sequences that suppress or amplify spontaneous oscillations. The second order model is given by equations 1–2. The following matrices describe the parameters of these equations:

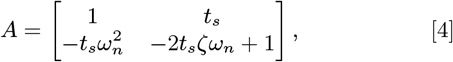

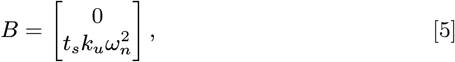

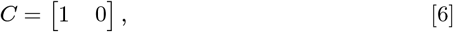

where *ω_n_* = 2*πf_n_* is the natural frequency (*rad/s*) of the evoked response, *ζ* is the damping ratio, *k_u_* is a scalar gain, and *t_s_* (s) is the sampling period.

We considered two distinct signals to characterize the spontaneous oscillations (*b*(*k*)) in the generalized neural dynamics model. The first signal type is given by a sinusoidal function of frequency *f_n_*:

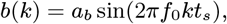

where *a_b_* is a constant that defines the signal amplitude. The second signal type is a sinusoidal function of frequency *f*_0_ whose amplitude envelope is modulated by a sinusoidal of lower frequency (*f_l_*). The following equation describes this signal:

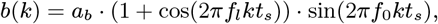

The nominal values of *f_n_* and *ζ* were set equal to 20 Hz and 0.092, respectively. These parameters correspond to the frequency and damping of the dominant mode in the patient evoked response model described below. Parameters *a_b_, f*_0_, and *f_l_* were equal to 10 *μ*V, 20 Hz and 2 Hz, respectively. The temporal and time-frequency responses to stimulation of the neural circuit’s synthetic model with the above nominal parameters are depicted in Figs. 2a-2b. To understand whether the stimulation pulse sequences derived with the nominal evoked response model differed from those one can obtain with distinct evoked response dynamics, we characterized how deviations in *ζ* and *f_n_* impact the optimized stimulation sequences. Parameter *ζ* was varied from 0.1 to 1, and *f_n_* from 10 to 30 Hz. The range used for the damping ratio spans the neural evoked dynamics from a lightly to a highly damped response We selected a natural frequency range that deviates +/- 50 % from the nominal natural frequency (*f*_0_=20 Hz) of the spontaneous oscillations.

**Fig. 2.**
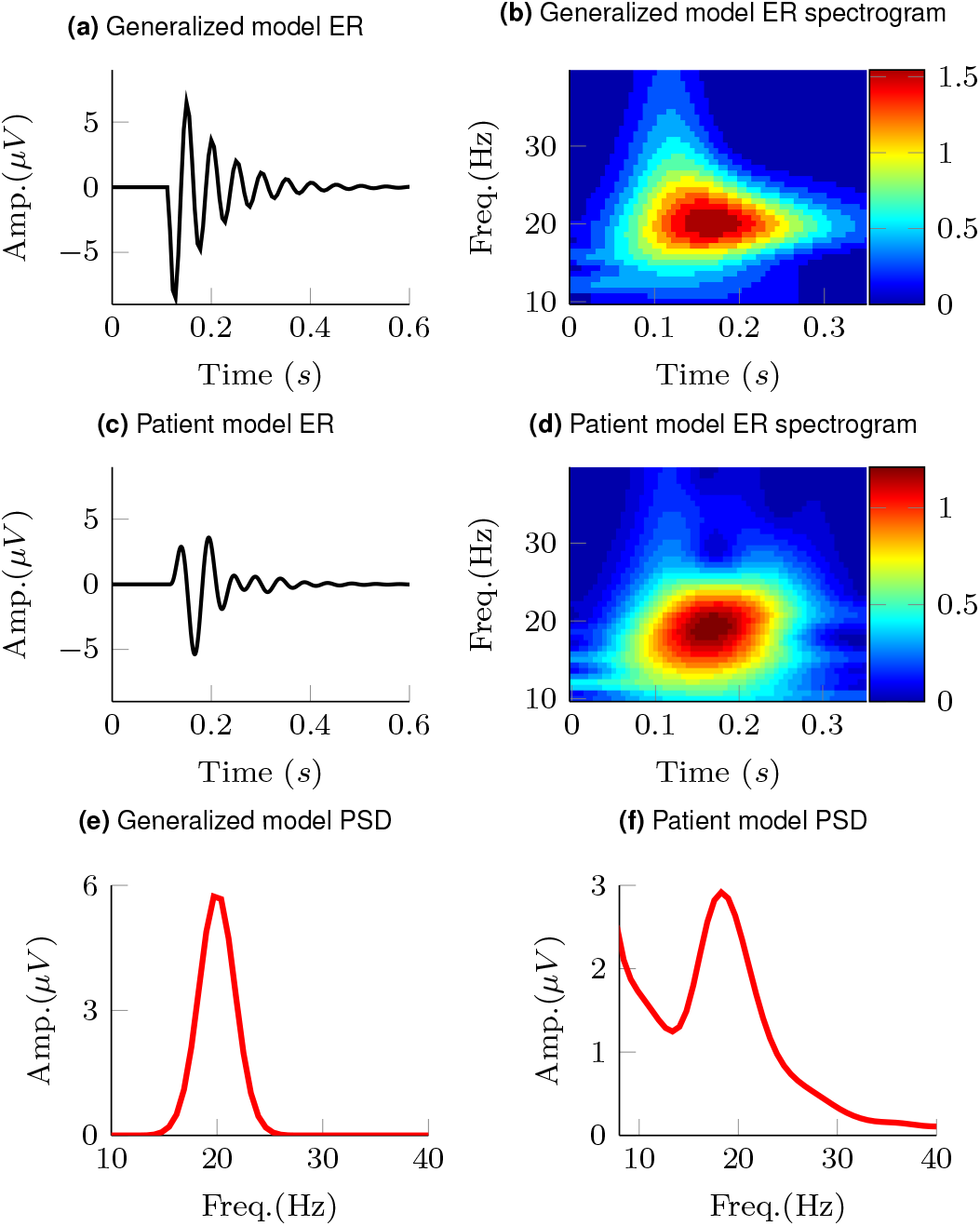
Stimulation-evoked responses for the generalized and patient-specific models are presented in **(a)** and **(c)**, respectively. The patient-specific model characterizes the neural response in the GPi evoked by stimulation in the GPi in a PD patient. Wavelet spectrograms (time-frequency maps) of the stimulation-evoked responses for the generalized mathematical model and patient-specific model are presented in **(b)** and **(d)**. Power spectral density (PSD) of spontaneous LFP activity for the generalized and patient-specific models are presented in **(e)** and **(f)**

#### Patient-specific model

A model of both spontaneous and stimulation-evoked neural activity in the GPi of a Parkinson’s disease patient was used to assess whether the results obtained with the generalized, low-order model can be applied to more complex and realistic neural dynamics. The patientspecific model was presented in a previous study in which the input-output relationship between stimulation in the GPi and evoked responses in the GPi was estimated using instrumental variable system identification (12). Briefly, the equations of differences describing the patient’s evoked response have two imaginary pairs of poles (modes) with natural frequencies equal to 14 and 20 Hz, and corresponding damping ratios equal to 0.22 and 0.092. The frequency response of the evoked response mathematical model indicated that the largest gain in the transfer function from stimulation input to evoked response is at approximately 20 Hz. See time-frequency response of the neural dynamics evoked by stimulation in Figs. 2c-2d and a more detailed description of the model in (12). Having evoked response dynamics with their largest gain at 20 Hz implies that periodic stimuli at 20 Hz result in the largest steady state, periodic evoked responses at this frequency. Therefore, stimulation pulses can be used to modulate 20 Hz neural activity with minimum amplitude.

The spontaneous neural activity used in the patient model was characterized with synthetic LFP signals that match the PSD observed in the GPi of a Parkinson’s disease patient in the resting, awake state. The patient evoked response model and PSD can be found in (12). The synthetic LFP signal is given by

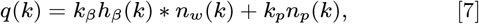

where * is the convolution operator, *h_β_* is a first order Butterworth filter with a pass-band in the 17-21 Hz range, *n_w_* is zero mean Gaussian white noise with unitary power, *n_p_* is a pink noise signal with unitary power, and *k_β_* = 12.6 and *k_p_* = 6 are constant scalars that are adjusted to match the PSD of the studied patient. The convolution between filter *h_β_* and the white noise *n_w_* is equal to the modeled beta band oscillations. The pink noise *n_p_* is used to characterize the 1/*f* background activity observed in LFP neural recordings (3, 4).

The signal targeted for modulation (*b*(*k*)) was obtained by filtering the LFP (*q*(*k*)) in the 14-24 Hz band using a second order Buttherworth filter. The power spectral density (PSD) of the synthetic LFP is shown in Fig. 2f.

### B. Stimulation constraints

We considered stimulation sequences subject to two distinct sets of constraints in our analysis and optimization algorithms. These stimulation constraints are described below.

#### 1. Stimulation pulses with adjustable amplitude

This class of stimulation sequences consists of pulses whose amplitude is freely selected by the optimization at any sample time. An example of a stimulation sequence with adjustable amplitude is shown in Fig. 3a and its corresponding evoked response is shown in Fig. 3b. For the mathematical formulation of our optimization problem, we employed the instantaneous normalized stimulation input 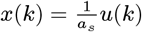 as the free optimization variable. This variable can take any value in the continuous compact interval [–1, 0]. *a_s_* is a constant equal to the maximum allowed stimulation current. The input *x*(*k*) = –1 corresponds to stimulation pulse with the largest allowed amplitude and *x*(*k*) = 0 corresponds to the stimulation current that does not evoke a neural response. The values of the stimulation variable are negative or zero because the negative phase of the stimulation pulse (cathodal stimulation) was assumed to evoke the neural response (11). It is noteworthy that using the same modeling framework but with a change in the saturation limits one could also characterize evoked responses dominated by the anodal phase of the stimulus.

**Fig. 3.**
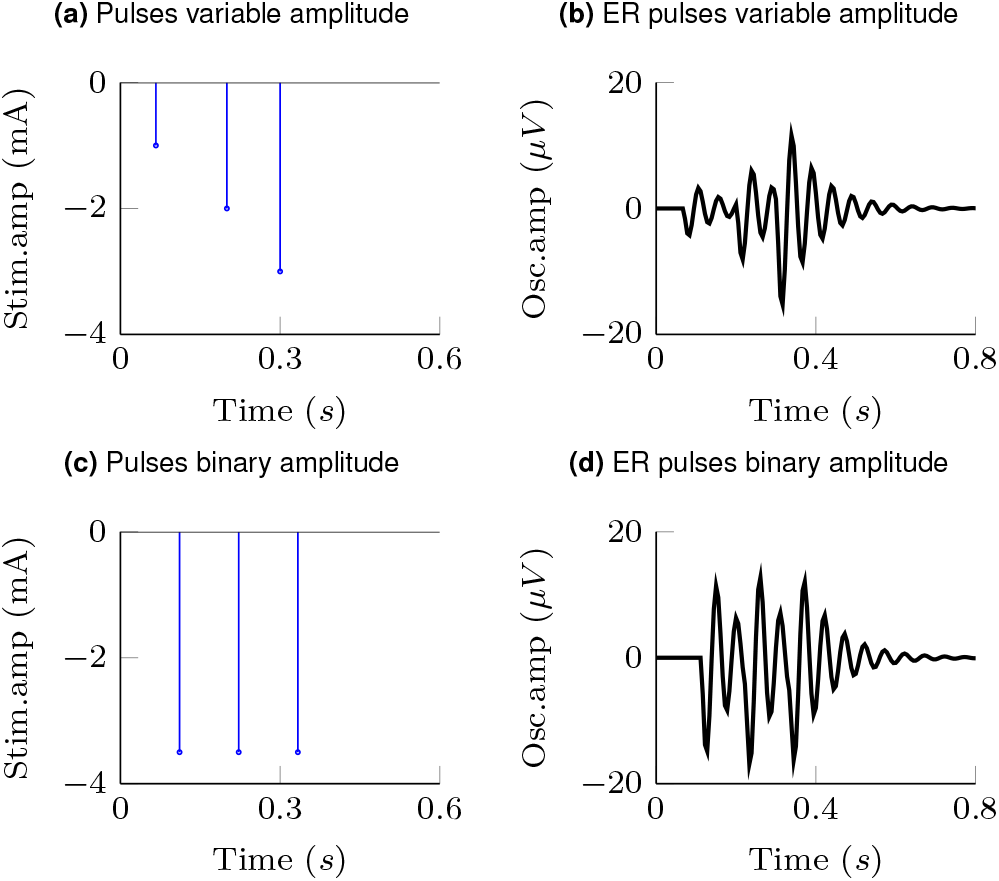
Examples of two different stimulation pulse patterns considered in this study and the corresponding evoked responses computed with the generalized, low-order model. A sequence of three pulses with adjustable amplitude and corresponding evoked response are presented in **(a)** and **(b)**, respectively. A sequence of pulses with binary amplitude values and corresponding evoked responses are shown in **(c)** and **(d)**.

#### 2. Stimulation pulses with binary amplitude values

The second class of stimulation sequences consists of pulses whose amplitude can be equal to either zero or a non-zero constant. This constraint on the stimulation pulse amplitude was studied only because present closed-loop neurostimulation systems allow us to deliver these sequences in humans (12), but not pulses whose amplitude can be adjusted in real-time. An example of a stimulation sequence with binary pulse amplitudes is shown in Fig. 3c and its corresponding evoked response is shown in Fig. 3d. We used the instantaneous normalized stimulation input 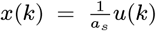 as the optimization variable. *x*(*k*) can take a value equal to either minus one or zero. Hence, *a_s_* is the stimulation pulse amplitude when *x*(*k*) = –1.

### C. Optimization approach

The goal of the optimization is to discover stimulation pulse sequences that maximize suppression or amplification of frequency-specific neural activity given the constraints described above. The mathematical function we aim to minimize is the 2-norm of the difference between the spontaneous neural activity and a reference signal to be tracked. This function is given by

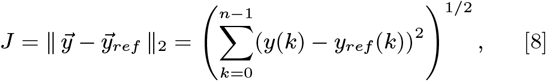

where || · ||_2_ represents the 2-norm and 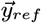 is the reference signal. The reference signal is set equal to zero when we want to suppress the spontaneous oscillations (i.e., 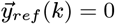 for all *k*). When we want to amplify the targeted neural activity, we set the reference signal equal to a sinusoidal with the same frequency and phase as the spontaneous oscillations but with a larger amplitude.

We considered electrical stimulation sequences with pulses delivered at samples *k* (i.e., samples at times *kt_s_*). A sampling period *t_s_* = 1/180 s was used in the optimization routines to ensure that stimulation pulses were generated with a frequency smaller than or equal to the sampling frequency (*f_s_* = 180 Hz). This frequency is smaller than the maximum frequency allowed in commercial DBS systems approved by the FDA for the treatment of neurological conditions (13). This sampling frequency is also large enough to capture the neural dynamics associated with spontaneous and stimulation-evoked oscillations with a frequency below 30 Hz. To formulate the optimization problem, we represent the solution to the equations of differences from expressions 1,2 using a matrix multiplication:

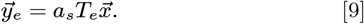

Vector 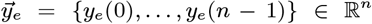 represents the evoked response at samples *k* = 0, … *n* – 1. Vector 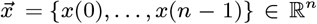 is the optimization variable and represents the normalized amplitude of the stimulation pulses. *T_e_* is a 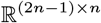 Toeplitz matrix that maps the stimulation sequences to the evoked response (14). The combined stimulation-evoked and spontanoues neural activity represented in vector form 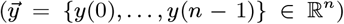 is given by

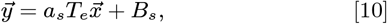

where 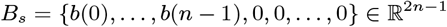.

The optimization problem for stimulation pulses with adjustable amplitude is described by the expression

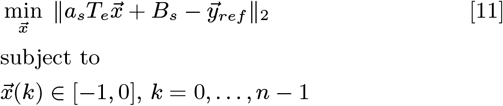

In the above optimization problem, *a_s_* is set equal to the maximum allowed stimulation current. This optimization problem is convex. Therefore, if a solution exists, it is unique and the global minimum (15). We denote the solution to this optimization problem as 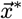.

For stimulation pulses with binary amplitude values, the optimization problem is

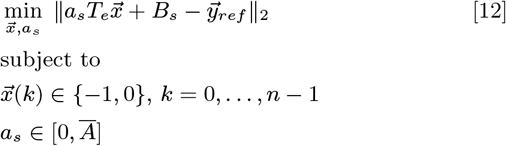

In this case, the constant scalar *a_s_* becomes an optimization variable. 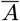 is the maximum value that *a_s_* is allowed to take (i.e., maximum stimulation current). The optimal values for this optimization problem are denoted as 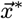 and 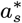. This optimization problem is not convex. Therefore, a minimum solution, if it exists, is not guaranteed to be unique or global (15). We used mixed-integer optimization algorithms to compute solutions to the optimization problem in expression 12.

#### Optimization algorithms and implementation

All the simulations and optimization algorithms were implemented in MATLAB (MathWorks, Natick, MA, USA) and a computer equipped with an Intel Core i7-8700K CPU processor (3.7 GHz) and a 32 GB RAM memory. For the optimization, we integrated the CVX package with Matlab (15) and solutions were obtained using the Gurobi solver. This solver was chosen because it has the capability of solving both convex and mixed-integer optimization problems (16, 17).

The optimization routines applied to both the low-order and patient-specific models were implemented with 2 s long LFP time series (*n* =360 samples). This period was sufficient to determine the steady-state stimulation patterns needed to modulate the targeted oscillations. It also enabled us to attain a solution to the mixed-integer optimization problem with a desired gap and within a reasonable time (<20 s). At each step, the solver internally calculates the best known objective function value for a feasible solution and its lower bound. The optimal objective function value is always between these two values. When the relative gap between these objective bounds is smaller than the “MIPGap” parameter of the solver, the optimization terminates. The MIPGap was set equal to 0.1 for our optimization.

### D. Assessment of neural modulation

We quantified the degree of modulation achieved by pulse sequences with adjustable or binary stimulation amplitudes using the optimal value of the optimization cost function 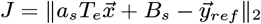.

## 3. Results

### A. Phase modulation of stimulation pulses mediate optimal suppression and amplification of oscillations with a constant amplitude envelope

The stimulation sequences derived with the nominal generalized model (nominal damping and natural frequency) that optimally suppressed or amplified spontaneous oscillations with a <u>constant amplitude</u> consist of pulse trains locked to a specific phase of the oscillations. Fig. 4 shows the wavelet spectrograms (time-frequency maps) of the modulated signal and the temporal evolution of the targeted oscillations, modulated signal, and stimulation pulses for the following scenarios: suppression of targeted oscillations using pulses with an adjustable amplitude (max. amplitude *a_s_*=2.5 mA), suppression of targeted oscillations using pulses with an adjustable amplitude (max. amplitude *a_s_*=0.4 mA), suppression of targeted oscillations using pulses with the binary amplitude constraint, and amplification of targeted oscillations using pulses with an adjustable amplitude (max. amplitude *a_s_*=2.5 mA). For brevity, we do not show the patterns for pulses with the binary amplitude constraint when amplification was the optimization goal.

**Fig. 4.**
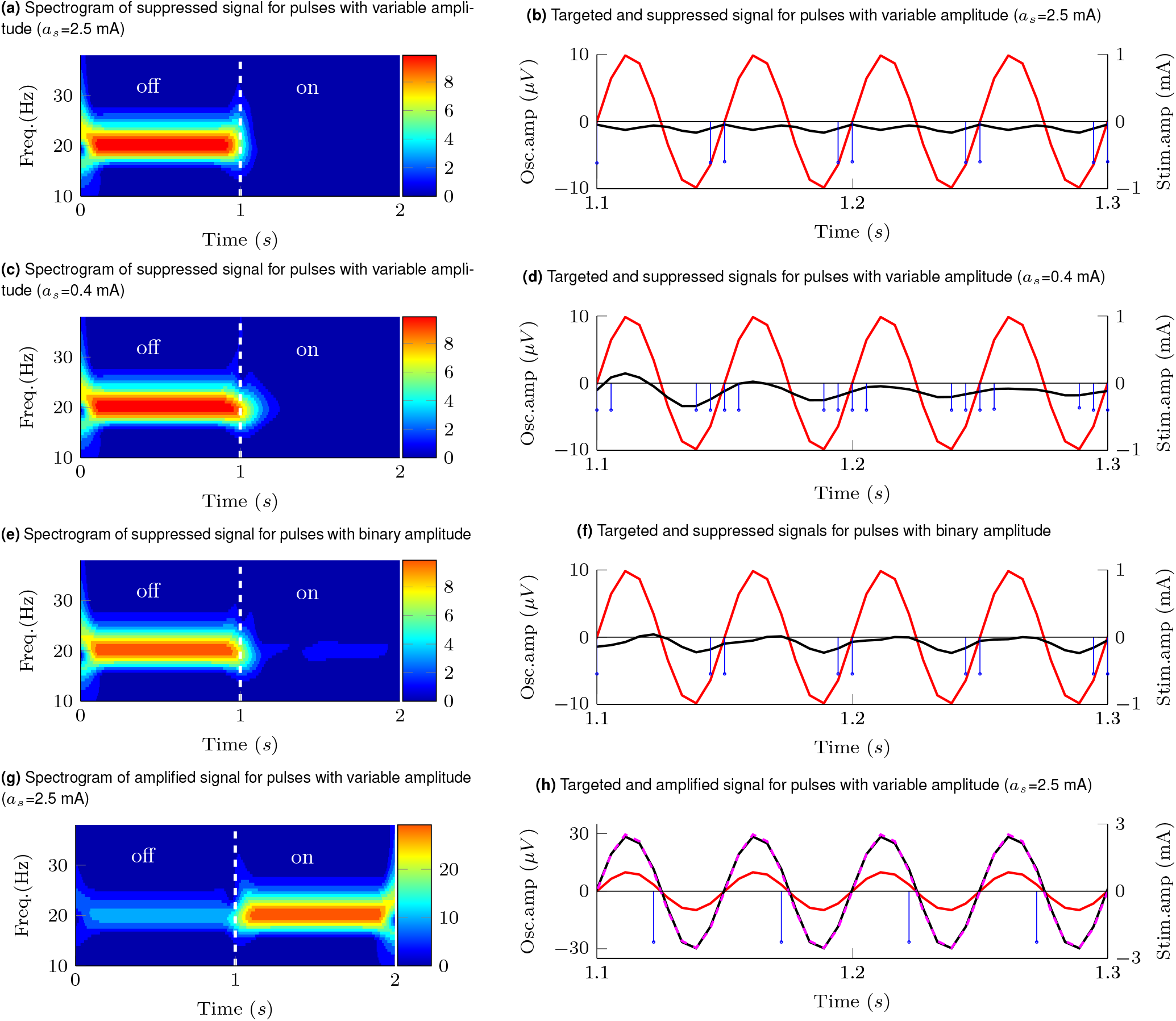
Effect of stimulation pulses on oscillations with constant amplitude envelope. Targeted oscillations 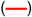, modulated (suppressed/amplified) oscillations (—), reference signal for amplification 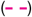, optimal stimulation sequence 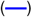 Targeted signal, modulated signal, and Wavelet spectrograms of the modulated signal during a transition from off- to on-stimulation are shown for sequences with the following characteristics: **(a,b)** pulses of adjustable amplitude (max. amplitude *a_s_*=2.5mA) that suppress the targeted oscillations, **(c,d)** pulses of adjustable amplitude (max. amplitude *a_s_*=0.4 mA) that suppress the targeted oscillations, **(e,f)** pulses with the binary amplitude constraint that suppress the targeted oscillations, **(g,h)** pulses of adjustable amplitude (max. amplitude *a_s_*=2.5 mA) that amplify the targeted oscillations.

The phase locking pattern was observed for pulses with both an adjustable amplitude and the binary amplitude constraint when the optimization goal was either suppression or amplification of the targeted oscillations. Nonetheless, the optimal cost for stimulation sequences with an adjustable amplitude (J=23.4 for suppression and J=49.3571 for amplification with *a_s_*=2.5 mA) is smaller than the cost for sequences with the binary amplitude constraint (J=410.2 for suppression and J= 687.986 for amplification).

We considered two constraints on the maximum stimulation amplitude (*a_s_*=2.5 mA and *a_s_*=0.4 mA) for the pulses with an adjustable amplitude. The relative effect of using a maximum current *a_s_*=2.5 mA vs. *a_s_*=0.4 mA helped us to understand how a decrease in the maximum allowed stimulation amplitude impacts the optimal pulse patterns. Constraints on the stimulation amplitude like those described above may be needed when the maximum electrical current cannot exceed a threshold in order to keep the electric charge within safe limits or when side effects due to stimulation need to be avoided. For targeted oscillations with a constant amplitude envelope, a decrease in the maximum allowed stimulation results in an increase in the number of pulses in each train aligned with the phase of the spontaneous oscillations. See spectrograms and pulse patterns in Figs. 4a and 4b vs. Figs. 4c and 4d. The greater number of pulses in each train results in an increase in the amplitude of the evoked responses (temporal summation) and thereby an enhanced ability to create constructive or destructive interference. It is worth mentioning here that the pulse frequency is limited by the sampling rate used in the mathematical models (i.e., 180 Hz).

### B. Pulse frequency, amplitude, and phase modulation mediate optimal suppression and amplification of oscillations with a time-varying amplitude envelope

The stimulation sequences that optimally suppress or amplify oscillations with a time-varying amplitude envelope exhibit a combination of phase, frequency, and amplitude pulse modulation for the case in which the pulse’s normalized amplitude is allowed to take any value in a continuous range between 0 and 1 (maximum current *a_s_* =2.5 mA). Pulse sequences with a higher frequency and amplitude are generated at specific phases of the spontaneous oscillations to increase the amplitude of the evoked responses and thereby achieve optimal destructive or constructive interference between the spontaneous and evoked oscillations. Figs. 5b and 5h show the pulse sequences that suppress and amplify the targeted oscillations (*a_s_* =2.5 mA), respectively.

**Fig. 5.**
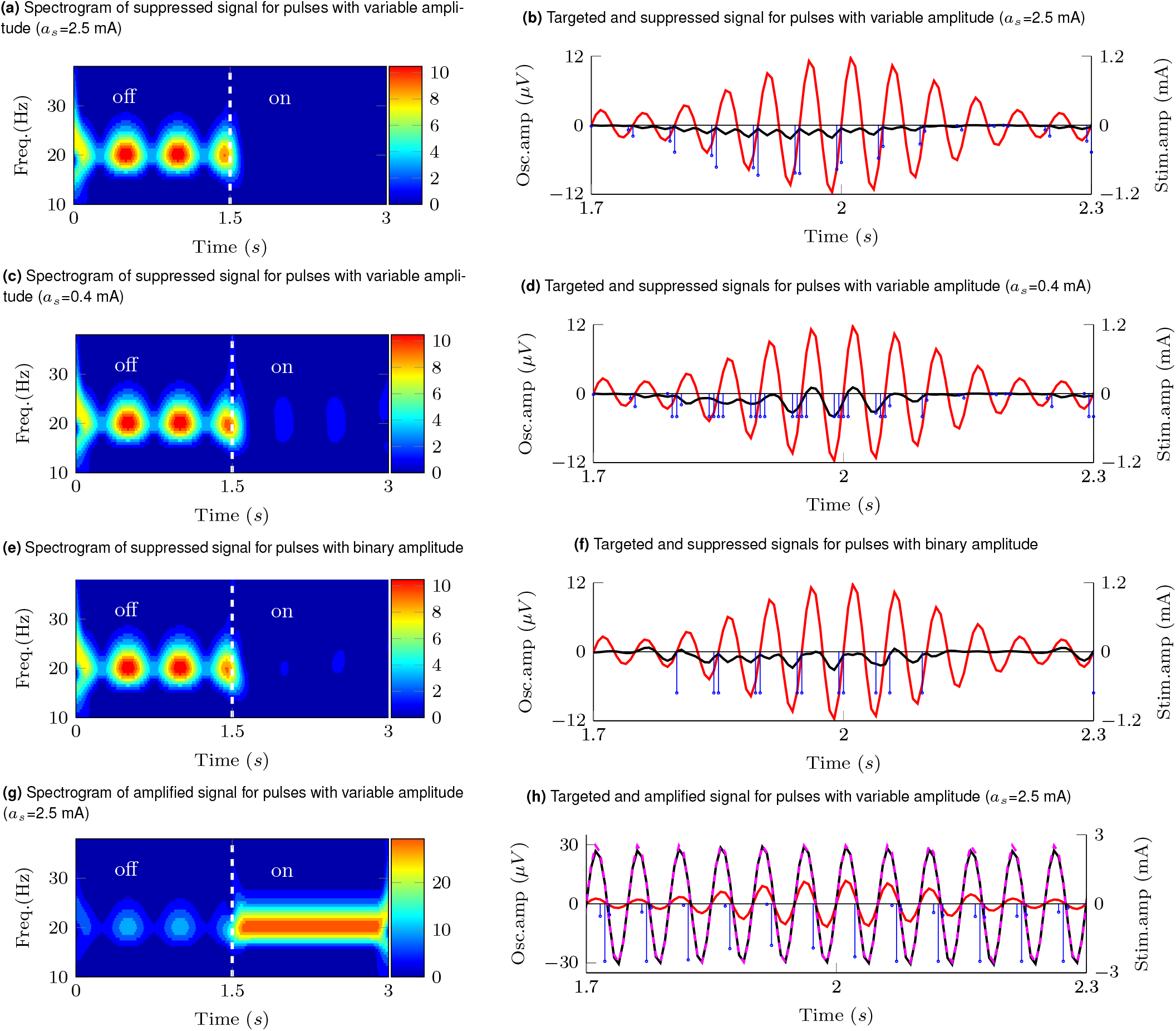
Effect of stimulation pulses on oscillations with time-varying amplitude envelope. Targeted oscillations 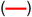, modulated (suppressed/amplified) oscillations (—), reference signal for amplification 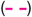, optimal stimulation sequence (—). Targeted signal, modulated signal, and wavelet spectrograms of the modulated signal during a transition from off- to on-stimulation are displayed for sequences with the following characteristics: **(a,b)** pulses of adjustable amplitude (max. amplitude *a_s_* = 2.5*mA*) that suppress the targeted oscillations, **(c,d)** pulses of adjustable amplitude (max. amplitude *a_s_* = 0.4*mA*) that suppress the targeted oscillations, **(e,f)** pulses of constant amplitude that suppress the targeted oscillations, **(g,h)** pulses of adjustable amplitude (max. amplitude *a_s_* = 2.5*mA*) that amplify the targeted oscillations.

For the pulses with an adjustable amplitude, our results indicate that pulse frequency modulation becomes more predominant than pulse amplitude modulation to maximize the suppression or amplification of the targeted oscillations as the value of the maximum allowed pulse amplitude (*a_s_*) is reduced from 2.5 to 0.4 mA. The spectrograms and pulse patterns in Figs. 5a and 5b vs. Figs. 5c and 5d illustrate how the reduction in the amplitude constraint influences the pulse patterns.

The solution to the optimization problem with the binary amplitude constraint on the pulses consists of sequences exhibiting a combination of pulse frequency and pulse phase modulation. When suppression was the optimization objective, these pulses were delivered only when the amplitude envelope of the oscillations was above a threshold. Fig. 5f vs. 5b show how stimulation is turned off when the envelope of the oscillations is below a threshold (time 2.1 s and after). This threshold-based strategy ensures that oscillations evoked by stimulation pulses with a fixed amplitude do not amplify the spontaneous oscillations when these oscillations are small compared to the evoked responses.

The optimization cost associated with the pulses with adjustable amplitude was smaller than the cost associated with the pulses with the binary amplitude constraint (J=22.7 vs. 353.8 for suppression and J=54.2147 vs. 912.569 for amplification). Therefore, pulse sequences with adjustable amplitude can achieve greater suppression or amplification than pulses with a binary amplitude constraint. The optimization results and cost function values for the studied pulse amplitude constraints and generalized model are summarized in Table 1.

**Table 1.** Summary of optimization results for the nominal generalized neural dynamics model.

|  | Oscillations with a constant amplitude envelope | Oscillations with a time-varying amplitude envelope |
| --- | --- | --- |
| Stimulation pulses with an adjustable amplitude (max. $a_s = 2.5$ mA) | Frequency, phase, and amplitude modulation (J=23.4 for suppression and J= 49.35 for amplification) | Frequency, phase, and amplitude modulation (J=22.7 for suppression and J=54.2147 for amplification) |
| Stimulation pulses with binary amplitude values (max. $a_s = 2.5$ mA) | Phase modulation (J=410.2 for suppression and J=687.986 for amplification) | Phase modulation (J=353.8 for suppression and J=912.569 for amplification) |

### C. Changes in the evoked response natural frequency and damping affect the degree of modulation but not the structure of the pulse sequences

The degree to which the optimized stimulation patterns modulate the targeted oscillations depends on the damping and natural frequency of the stimulation-evoked responses (Fig. 6). The optimal value of the cost function (*J*) is a monotonically increasing function of the damping ratio for both targeted oscillations with constant and time-varying envelope (Figs. 6a, 6b, 6d, and 6e). When the natural frequency of the evoked response matches the frequency of the targeted oscillations, the lowest value of the optimal cost function is attained. The optimal cost increases as the natural frequency moves away from the targeted oscillations’ frequency (Figs. 6a, 6c, 6d, and 6f).

**Fig. 6.**
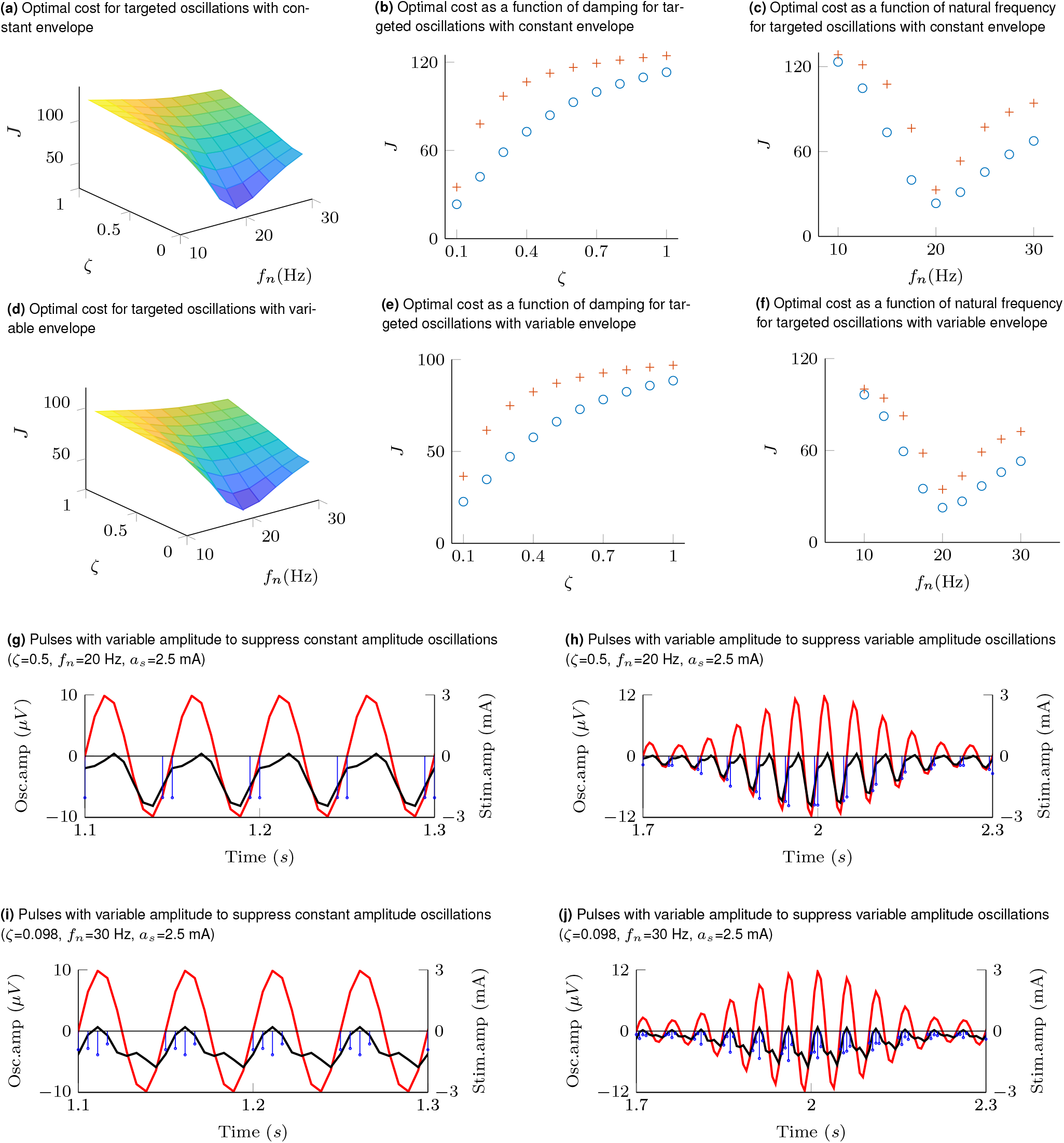
The surface plots in **(a)** and **(d)** illustrate the optimal cost function value (*J*) as a function of the damping ratio (*ζ*) and natural frequency (*f_n_*) when the targeted oscillations are suppressed via pulse sequences with adjustable amplitude (*a_s_* =2.5 mA). Targeted oscillations with a constant envelope are considered in **(a)**, and with a time-varying envelope in **(d)**. The curves in **(b)** and **(e)** show the optimal value of *J* as a function of *ζ* when *f_n_*=20 Hz (nominal value) for targeted oscillations with a constant and time-varying envelope, respectively. These curves illustrate the optimal cost function values for pulse sequences with adjustable amplitude whose maximum allowed amplitude (*a_s_*) was set equal to 2.5 mA 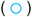, and 0.4 mA 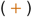. The curves in **(c)** and **(g)** show the optimal value of *J* as a function of *f_n_* when *ζ*=0.092 (nominal value) for targeted oscillations with a constant and time-varying envelope, respectively. Examples of pulse sequences and modulated signals for evoked responses with a damping ratio *ζ* =0.5 (different from its nominal value) and *f_n_* =20 Hz (nominal) are shown in **(g)** and **(h)** for targeted oscillations with a constant and time-varying envelope, respectively. The plots in **(i)** and **(j)** show examples of pulse sequences and modulated signals for evoked responses with *ζ* =0.092 (nominal) and *f_n_* =30 Hz (different from nominal) when the targeted oscillations have a constant and time-varying amplitude envelope, respectively.

The structure of the pulse patterns (i.e., pulse phase, amplitude, and frequency modulation) did not change when the damping or natural frequency were varied. Figs. 6g and 6h illustrate examples of optimal pulse sequences with adjustable amplitude for evoked responses with a damping ratio equal to 0.5 and natural frequency equal to 20 Hz. While this damping ratio significantly departs from the nominal value (0.092), pulse phase modulation is preserved to suppress the targeted oscillations with constant amplitude, and a combination of phase, amplitude, and frequency modulations is preserved to suppress the targeted oscillations with time-varying amplitude. See Figs. 6g vs. 4b and 6h vs. 5b. Figs 6i and 6j show that when the nominal natural frequency (20 Hz) is increased to 30 Hz, the pulse phase modulation is preserved to suppress the targeted oscillations with constant amplitude, and the combination of phase, amplitude, and frequency modulations is preserved to suppress the targeted oscillations with time-varying amplitude. See Figs. 6i vs. 4b and 6j vs. 5b.

### D. Stimulation sequences for patient-specific neural dynamics model matched those obtained with the generalized model

Stimulation pulse sequences with adjustable amplitude employed frequency, phase, and amplitude pulse modulation to attain maximum suppression or amplification of targeted oscillations in the patient-specific model of the GPi neural dynamics (Figs. 7a and 7b). As we reduced the maximum allowed stimulation amplitude from 2.5 mA to 1.25 mA, frequency modulation rather than amplitude modulation dominated the pulse patterns to achieve maximum suppression (Figs. 7c and 7d). Stimulation pulses with the binary amplitude constraint used phase and frequency modulation to modulate the targeted oscillations (Figs. 7e and 7f). The reduction in the 2-norm of the modulated signal was greater for the adjustable amplitude pulses (J=23.4 for suppression and J=49.3571 for amplification) than for the binary amplitude pulses (J=410.2 for suppression and J= 687.986 for amplification). These results agree with those obtained with the generalized neural dynamics model.

**Fig. 7.**
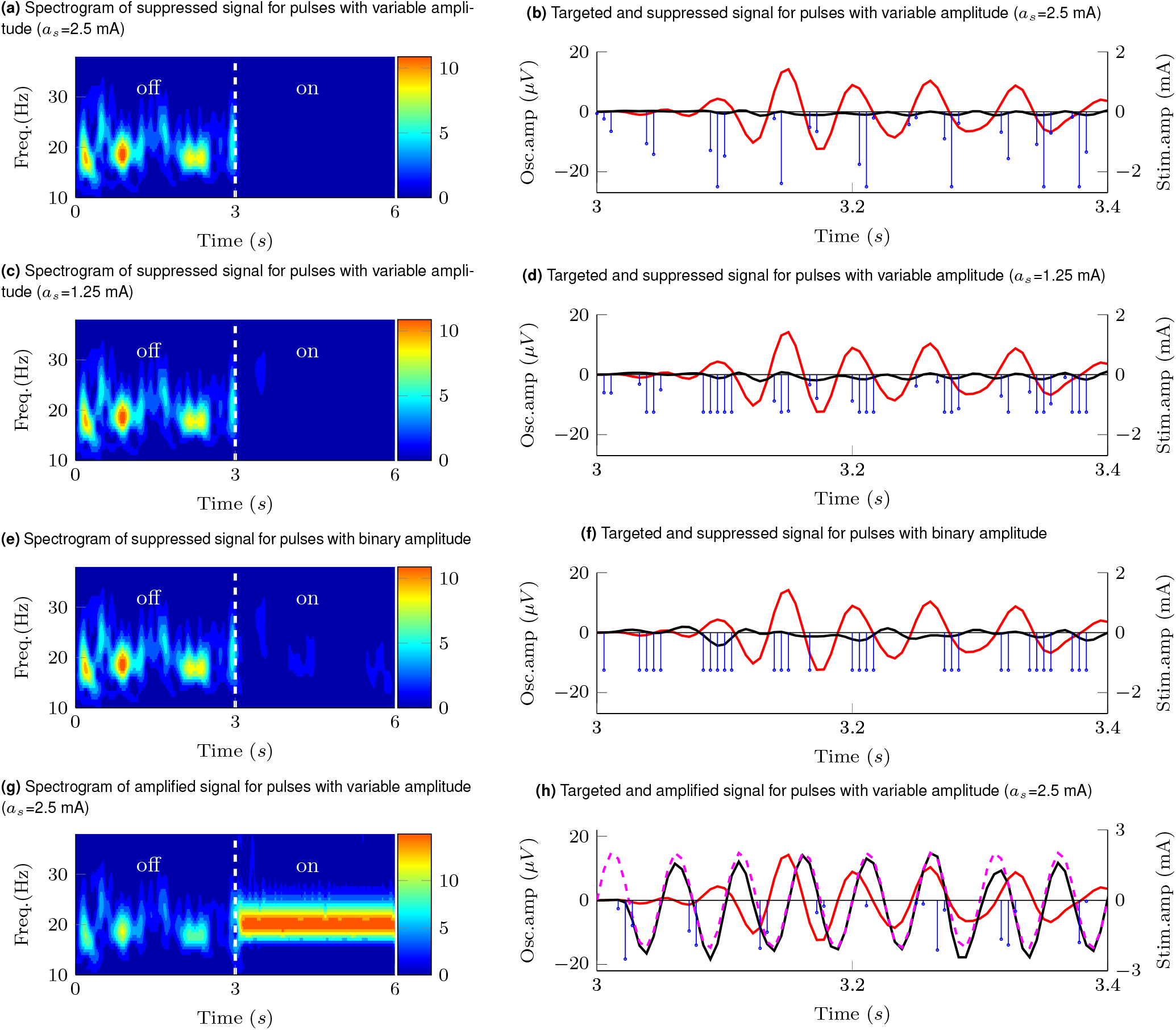
Effect of stimulation pulses on patient-specific GPi neural dynamics. Targeted oscillations 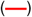, modulated (suppressed/amplified) oscillations (—), reference signal for amplification 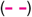, optimal stimulation sequence 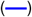. Targeted signal, modulated signal, and wavelet spectrograms of the modulated signal during a transition from off- to on-stimulation for sequences with the following characteristics: **(a,b)** pulses of adjustable amplitude (max. amplitude *a_s_* =2.5mA) that suppress the targeted oscillations, **(c,d)** pulses of adjustable amplitude (max. amplitude *a_s_* =1.25 mA) that suppress the targeted oscillations, **(e,f)** pulses of constant amplitude that suppress the targeted oscillations, **(g,h)** pulses of adjustable amplitude (max. amplitude *a_s_* =2.5 mA) that amplify the targeted oscillations.

Fig. 8 shows the PSD curves of the modulated signal when the optimization goal was to suppress or amplify the targeted neural activity. These PSD curves show that stimulation pulses with both adjustable and binary amplitudes can effectively modulate the mean power of frequency specific neural oscillations in the targeted frequency band (14-24Hz). They also indicate that the mean power reduction attained via pulses with adjustable amplitude is comparable to the reduction achieved via pulses with the binary amplitude constraint, even though the difference in the cost function values was more notable.

**Fig. 8.**
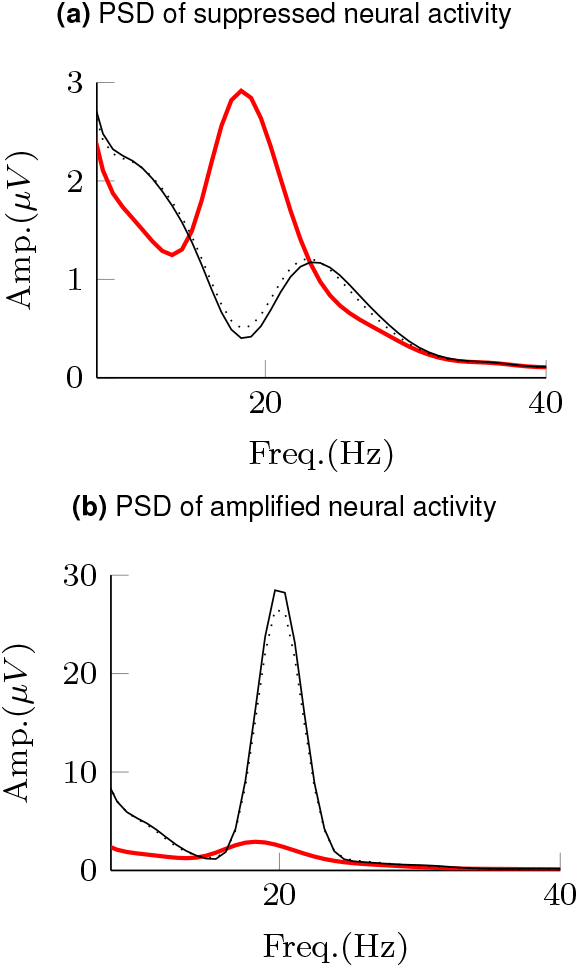
Power spectral density (PSD) curves of neural activity in the internal segment of the globus pallidus from a patient-specific mathematical model. PSD curves shown in the figure are associated with periods in the off-stimulation condition 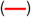 and when stimulation pulses with adjustable amplitude (–) and the binary amplitude constraint (⋯) were delivered. **(a)** and **(b)** show PSD curves during periods of suppression and amplification of frequency-specific neural activity, respectively.

## 4. Discussion

In this study, we identified electrical pulse patterns to effectively modulate spontaneous neural oscillations using stimulation-evoked neural responses. We leveraged mathematical optimization tools to discover pulse patterns that minimize (or maximize via reference tracking) the 2-norm of frequency-specific neural oscillations. To identify these pulse patterns, we employed mathematical models of spontaneous and stimulation-evoked oscillations. We considered stimulation-evoked response models with distinct natural frequency and damping coefficient as well as a subject-specific GPi model of a Parkinson’s disease patient. The distinct neural dynamics studied here enabled us to assess whether optimized pulse sequences preserve their structure. Our analysis revealed that, across the studied neural dynamics, a combination of phase, amplitude, and frequency modulation of the electrical pulses results in optimal suppression or amplification of the targeted oscillations. These pulse modulation strategies provide the rationale for the development of future closed-loop neuromodulation systems that employ stimulation-evoked responses and pulse modulation strategies to effectively control neural activity in real-time. These closed-loop neurostimulation systems will, in turn, be key to characterize the role of neural activity in brain function and ultimately advance personalized neuromodulations therapies.

### A. Mechanisms underlying optimized neural modulation

The optimized stimulation sequences identified in this study take advantage of phase, frequency and amplitude pulse modulation to achieve optimal suppression or amplification of targeted oscillations. Phase modulation (or time alignment) of the pulse trains relative to the targeted oscillations’ phase is required to effectively achieve either destructive or constructive interference between the spontaneous and evoked oscillations. We showed that when the amplitude envelope of the spontaneous oscillations is constant over time, phase modulation (or locking) is sufficient to achieve optimal suppression or amplification of the targeted oscillations. However, the envelope of real in-vivo spontaneous oscillations is not constant over time (1–4, 11, 12). Our results show that adjustments in the stimulation pulse frequency and amplitude are needed to create precise destructive or constructive interference between stimulation-evoked responses and spontaneous oscillations whose amplitude envelope varies over time. Pulse frequency modulation results in changes in the evoked response amplitude because of the principle of superposition (or temporal summation) observed in responses to stimulation trains (18–20). For this superposition or temporal summation to be effective, the intra-burst period of the pulse trains must be greater than the refractory period of the neuronal population being stimulated with the electrical pulses. If the inter-pulse period is smaller than the refractory period of the neural circuitry being activated, a temporal summation of evoked responses is not expected to occur (19, 21, 22). Pulse amplitude modulation can be employed, in addition or instead of frequency modulation, to adjust the evoked responses’ amplitude and suppress the spontaneous oscillations. Because the pulse amplitude (voltage or current) is limited in FDA-cleared or -approved neurostimulation devices to minimize the likelihood of tissue damage, we studied the effect of constraining the pulse amplitude on the pulse patterns. Our results indicate that when the maximum allowed pulse amplitude (upper bound) is reduced, frequency pulse modulation becomes more predominant than amplitude pulse modulation in order to achieve optimal suppression or amplification of the targeted oscillations. This trade-off is important for the implementation of closed-loop neuromodulation systems since they can use a combination of amplitude and frequency pulse modulation to attain a desired modulation performance given the device constraints and safety envelope.

### B. Generalization of results

We employed a generalized, low-order mathematical model of stimulation-evoked neural dynamics to characterize optimal pulse sequences that suppress (or amplify) spontaneous neural oscillations. The low-order model was characterized by a single natural frequency and damping coefficient. We studied the effect of varying these two parameters on the optimized pulse sequences to understand how invariant are the pulse modulation strategies across distinct neural dynamics. We also used a model of spontaneous and stimulation-evoked neural activity in the GPi of a Parkinson’s disease patient to identify the optimal pulse sequences in a more realistic scenario. Our results indicate that the structure of the optimal pulse patterns (phase, amplitude, and frequency pulse modulation) did not change when the stimulation-evoked neural dynamics were varied or when using the patient-specific model. This consistency across models indicates that pulse modulation strategies can be applied to suppress or amplify neural activity in distinct brain circuits and conditions, whenever spontaneous and stimulation-evoked responses of comparable magnitude are generated in the targeted neuronal population. For example, underdamped neural responses in the hippocampus CA1 area evoked by stimulation of the endopiriform nucleus (23) could be employed, together with our pulse modulation strategies, to control hippocampal, frequency-specific neural activity.

### C. From optimal pulse sequences to real-time closed-loop neuromodulation

How to embed the optimal pulse sequences studied in this article onto closed-loop stimulation devices is yet to be investigated. A foreseeable approach is to take the pulse sequences from a subject-specific model and embed them onto a closed-loop control system that adjusts the phase, amplitude, and frequency of the pulses based on realtime measurements of the oscillations’ amplitude and phase. A major challenge to implement this approach is that pulse sequences calculated using the proposed optimization are obtained based on knowledge of all the spontaneous oscillations samples (from *k* = 0 to *k* = *n*). Therefore, pulses delivered at time *k* depend on future values (*k* + 1, *k* + 2, …) of the spontaneous oscillations. This strategy is not possible to implement in a real-time system unless a prediction of the future spontaneous neural activity is available. Delivering the optimized pulse sequences based on present and past values of the modulated neural activity (i.e., delayed version of the optimal pulse sequence) may provide greater neuromodulation performance than phase-locked stimulation alone, but likely a reduced performance compared to the optimal stimulation sequences identified in this study because of the time delay introduced in the feedback loop. Future studies need to assess the performance of closed-loop systems with phase, amplitude, and frequency pulse modulation in order to optimize neural control technology. These technology advancements will be key to characterize how controlled changes in neural activity relate to brain function in the healthy and disease states, and to develop therapies tailored to each patient.

### D. Limitations

To characterize the optimal pulse modulation sequences, we used models of the stimulation evoked responses that consist of a linear dynamical system with stimulation inputs constrained to be square pulses. This model does not consider limits in the amplitude of the evoked and spontaneous neural oscillations and other nonlinearities present in neural circuits. Not including the nonlinear dynamics into the mathematical models can lead to inaccuracies in the pulse sequences calculated via the optimization routines. Nevertheless, our results are valid when considering neural dynamics operating in the linear region. Additionally, introducing a nonlinear element into the mathematical model of the evoked and spontaneous oscillations can result in a non-convex optimization problem whose solution is not guaranteed to be global. This study is also limited to stimulation sequences optimal when measuring the 2-norm of the targeted oscillations. We selected the 2-norm because it enables us to formulate a convex optimization problem and is intuitive mathematically. Yet, other norms (e.g., 1-norm and ∞-norm) and other cost functions may be more relevant for particular problems in neuroscience and to test specific hypotheses. For example, one may want to use the ∞-norm to determine the pulse sequences needed to minimize the maximum amplitude that oscillations can attain.

## ACKNOWLEDGMENTS

Research reported in this publication was funded by the Wallin Discovery Fund, the Engdahl Family Foundation, the National Institute of Neurological Disorders and Stroke (P50-NS123109, R01-NS037019, R37-NS077657), the University of Minnesota’s MnDRIVE (Minnesota’s Discovery, Research and Innovation Economy) Initiative-Postdoctoral Neuromodulation Fellowship granted to Hafsa Farooqi, and seed funds provided to David Escobar by the Department of Biomedical Engineering (Lerner Research Institute) and Neurological Institute at Cleveland Clinic.

